# Response of soil bacteria to PUREX chemicals suggests biomarker utility and bioremediation potential

**DOI:** 10.1101/2023.10.06.561230

**Authors:** Justin C Podowski, Sara Forrester, Dionysios A. Antonopoulos, Jennifer L. Steeb, Angela D. Kent, James J Davis, Daniel S. Schabacker

## Abstract

Chemicals involved in plutonium uranium reduction extraction (PUREX) have the potential to be released from nuclear reprocessing facilities and accumulate in the environment. In order to understand how soil microbial communities respond to contamination by PUREX chemicals, we carried out a series of microcosm experiments, exposing chemically diverse soils to a range of concentrations of key chemicals used in the PUREX process. We tested 4 PUREX chemicals, and 5 soil types using 16S rRNA amplicon sequencing, determining that responses of microbial communities are dependent on the soil type in which they reside, and that tributyl phosphate exposure appears to generate the most reproducible and detectable shifts in microbial communities. We identified a number of key taxa that are consistently enriched in soils exposed to tributyl phosphate. These key taxa are either in the family *Rhizobiaceae* or genus *Pseudomonas*. The relative abundance of these key taxa is concentration dependent, and their abundance remains elevated at least 100 days post initial exposure. Using whole-shotgun metagenomic sequencing, we reconstructed the genomes of these key taxa and find a number of putative phosphotriesterase genes found only in *Rhizobiaceae*. We find the abundance of phosphotriesterase genes is significantly higher in samples exposed to tributyl phosphate. These phosphotriesterase genes, which degrade tributyl phosphate into dibutyl phosphate and butanol, may serve as effective biomarkers for tributyl phosphate contaminated soil, as well as a method for future bioremediation.

**Importance:** Nuclear materials reprocessing facilities have the capacity to release toxic chemicals during normal operations or accidents. This study examines the ways in which chemicals involved with nuclear materials reprocessing impact microorganisms in the soil. Our intention was to understand the consequences of the release of these chemicals on ecosystems that may surround these reprocessing facilities. We find soil microbial communities change in response to some chemicals but not others, and that tributyl phosphate appears to generate the most reproducible and detectable shifts in microbial communities. Microorganisms in the family *Rhizobiaceae* increase in abundance in response to the addition of tributyl phosphate, and an examination of the genomes of these microbes suggest they may be able to break down tributyl phosphate to access the phosphosphate present in this chemical. Overall, this work demonstrates that changes in soil microbial communities in response to contamination with chemicals from nuclear materials reprocessing facilities may be predictable, and these responses could be leveraged to remediate contamination.

## Introduction

Accidental releases of waste and byproducts from nuclear materials processing facilities pose a grave danger to the health of ecosystems surrounding these facilities. Releases can take the form of dramatic and catastrophic events such as the Chernobyl and Fukushima disasters. But more commonly, these releases happen over long periods of time, as was the case for the Savannah River Site in South Carolina (1), sites around Oak Ridge National Laboratory and the Y-12 plant in Tennessee (2), and the Hanford site in Washington (3). While accumulation of radionuclides and heavy metals in the surrounding environment is often the focus of research and remediation, nuclear materials processing facilities also have the potential to release chemicals which have been less well studied, but that may have just as large of an impact on ecosystems. This is especially true for chemicals that may be easily aerosolized, be deposited, and accumulate far from a facility. These acids, organophosphates, hydrocarbons and more (4) have the potential to accumulate in ecosystems and cause great harm.

Plutonium uranium reduction extraction (PUREX) is the original method for purifying plutonium and uranium (5) from spent nuclear fuel and was used at the Hanford site (6) and Oak Ridge National Laboratory (7). Several chemicals involved in the PUREX process have been identified as the most likely to be released from nuclear materials processing plants during normal operation, due to their vapor pressure and volatility (4). These chemicals can be released through off gassing from cooling stacks, through venting of storage tanks, or from breakdown of storage tanks (8). In fact, hydrocarbons likely derived from storage tank leakage have been identified in soils surrounding the Hanford site (9).

Microorganisms in terrestrial environments not only comprise almost half of all belowground biomass (10) but also provide a wide array of ecosystem services (11) and changes in terrestrial microbial communities can act as predictors of changes in soil health (12). While microbial communities are to some degree functionally redundant, shifts in microbial diversity and taxonomy can have impacts on the ecosystem services they provide (13,14). As a result, impacts on terrestrial microbial communities by the release of chemicals from nuclear materials processing facilities have the potential to propagate to other parts of the terrestrial ecosystem.

Even though terrestrial ecosystems may be greatly impacted by chemical releases, terrestrial microbes may also serve as the greatest resource for remediation of contamination from nuclear facilities. Soil microbes carry out redox reactions that immobilize radionuclides (15,16), degrade complex organic solvents (17), and organophosphates (18). But not all microbes are created equal in this respect. Taxonomic groups such as *Rhizobia* (19) or *Pseudomonas* (20,21) are known to have the ability to break down a broad spectrum of chemicals. As such, it can be expected that the ability of some but not all microbes to degrade chemicals may lead to an enrichment of key microbes in response to accumulation of chemicals.

In an attempt to better understand the impacts of potential chemical releases from nuclear materials reprocessing facilities on terrestrial microbial communities, we carried out a series of laboratory microcosm experiments. We focused on key semi volatile chemicals used in plutonium uranium reduction extraction (PUREX) because they are relatively environmentally stable and tend to be adsorbed on soils and surfaces near the source facility.

## Results

### Community-level and individual ASV response to PUREX chemicals

The soils used in this study were selected to represent a wide array of chemical and physical properties. Overall, 5 unique soil types were used (Table 1, Tables S1-S3), with organic carbon percentages varying from 1.9 % to 16.7%, and cation exchange capacities varying from 10.7 meq/100g to 19.4 meq/100g. Soils were exposed to 4 PUREX chemicals, kerosene (K), kerosene with tributyl phosphate (KT), organic layer of kerosene with tributyl phosphate and nitric acid (KTO), and aqueous layer of kerosene with tributyl phosphate and nitric acid (KTA).

**Table 1.** Field history, soil texture and chemistry information for soils collected for each experiment. Each experiment involved a new collection of soil, so repeat chemistry measurements were performed.

| Soil ID | Experiment | Soil Textural Class | Organic Matter % | pH | % Sand | % Silt | % Clay | Cation Exchange Capacity (meq/100g) |
| --- | --- | --- | --- | --- | --- | --- | --- | --- |
| P02 | 1 | Loam | 8.6 | 6.8 | 43 | 44 | 13 | 19.0 |
| P15 | 1 | Loam | 4.6 | 6.7 | 35 | 49 | 16 | 13.2 |
| DAN | 1 | Loam | 8.8 | 6.7 | 46 | 47 | 7 | 18.3 |
| SAN | 1 | Sand | 1.9 | 7.6 | 91 | 6 | 3 | 14.1 |
| P02 | 2 | Sandy Loam | 9.1 | 6.8 | 53 | 34 | 13 | 15.8 |
| P15 | 2 | Loam | 4.9 | 7.2 | 41 | 48 | 11 | 11.6 |
| DAN | 2 | Loam | 3.7 | 7.9 | 35 | 41 | 24 | 17.4 |
| DRU | 2 | Loam | 9.6 | 7.0 | 39 | 39 | 22 | 18.7 |
| P02 | 3 | Sandy Loam | 9.2 | 6.6 | 62 | 28 | 10 | 17.7 |
| P15 | 3 | Loam | 5.5 | 6.8 | 46 | 38 | 16 | 10.7 |
| DAN | 3 | Sandy Clay Loam | 2.4 | 7.8 | 54 | 26 | 20 | 19.0 |
| DRU | 3 | Loam | 16.7 | 6.6 | 34 | 42 | 24 | 19.4 |

First, we exposed each soil type to each of the four chemicals at a concentration of 100 μl/g, and measured the community composition at 0-, 1-, 7- and 28-days following exposure. We observed changes in community composition in response to the KT, KTO, and KTA exposures (Figure 1). This response is not universal, and communities from soil types with higher sand content do not respond as strongly to any chemical amendments (Table 1, Figure 1). Overall, we observed the most dramatic response to the KTA condition. However, decreases in Shannon’s Diversity Index for KTA exposed samples suggest that these apparent differences are due to decreases in diversity rather than shifts in community composition (Figure 2).

**Figure 1:**
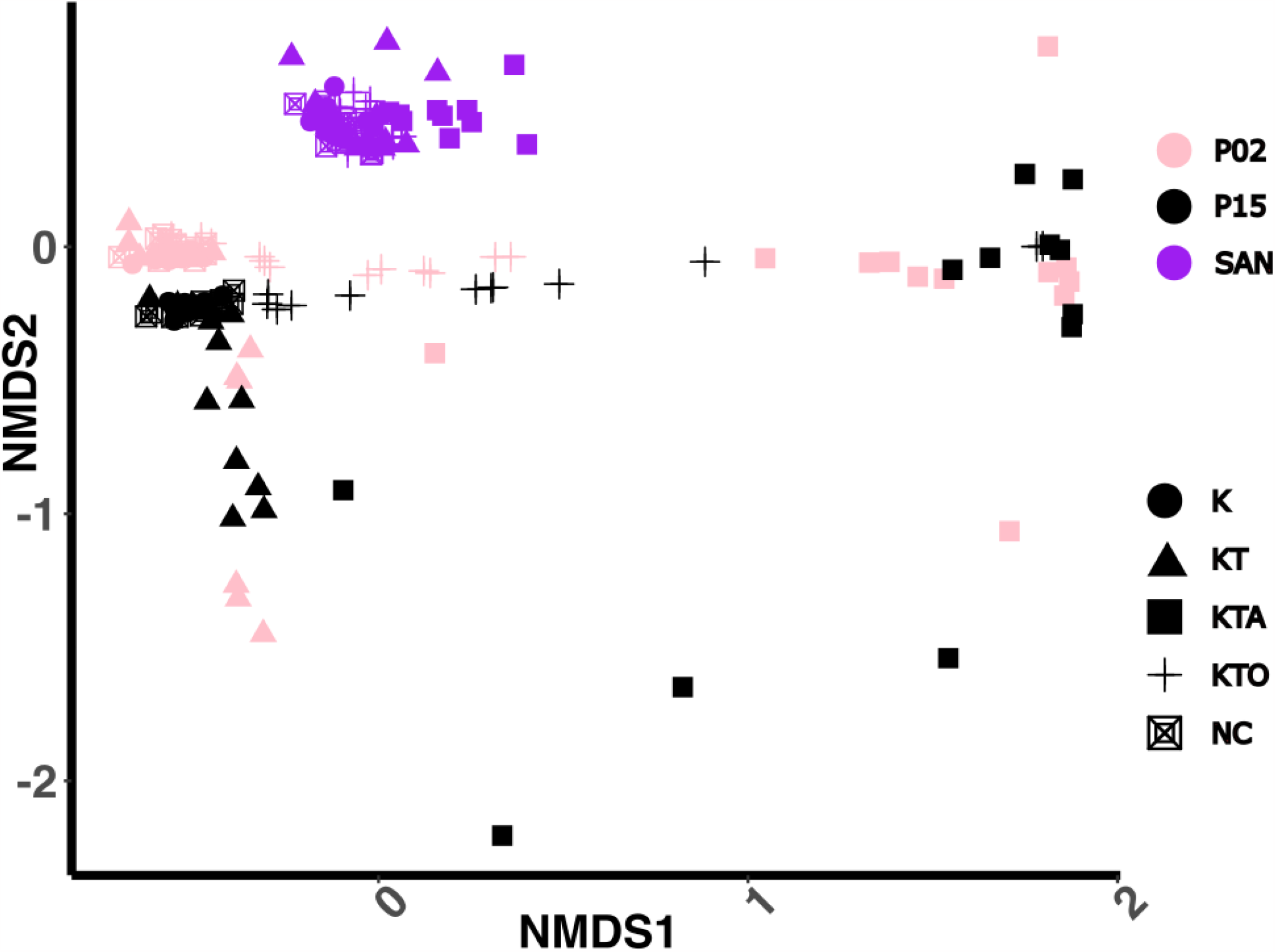
Non-metric Multi-dimensional Scaling (NMDS) ordination plot of Bray-Curtis Dissimilarity between 16S samples were extracted from the soil at 0, 1, 7 and 28 days following a 100 μl/g chemical exposure to each chemical. Plot points are colored based on soil type, and plot symbols are based on the chemical condition. Soil types are described in Table 1. Samples are from Experiment 1 (Table S1).

**Figure 2.**
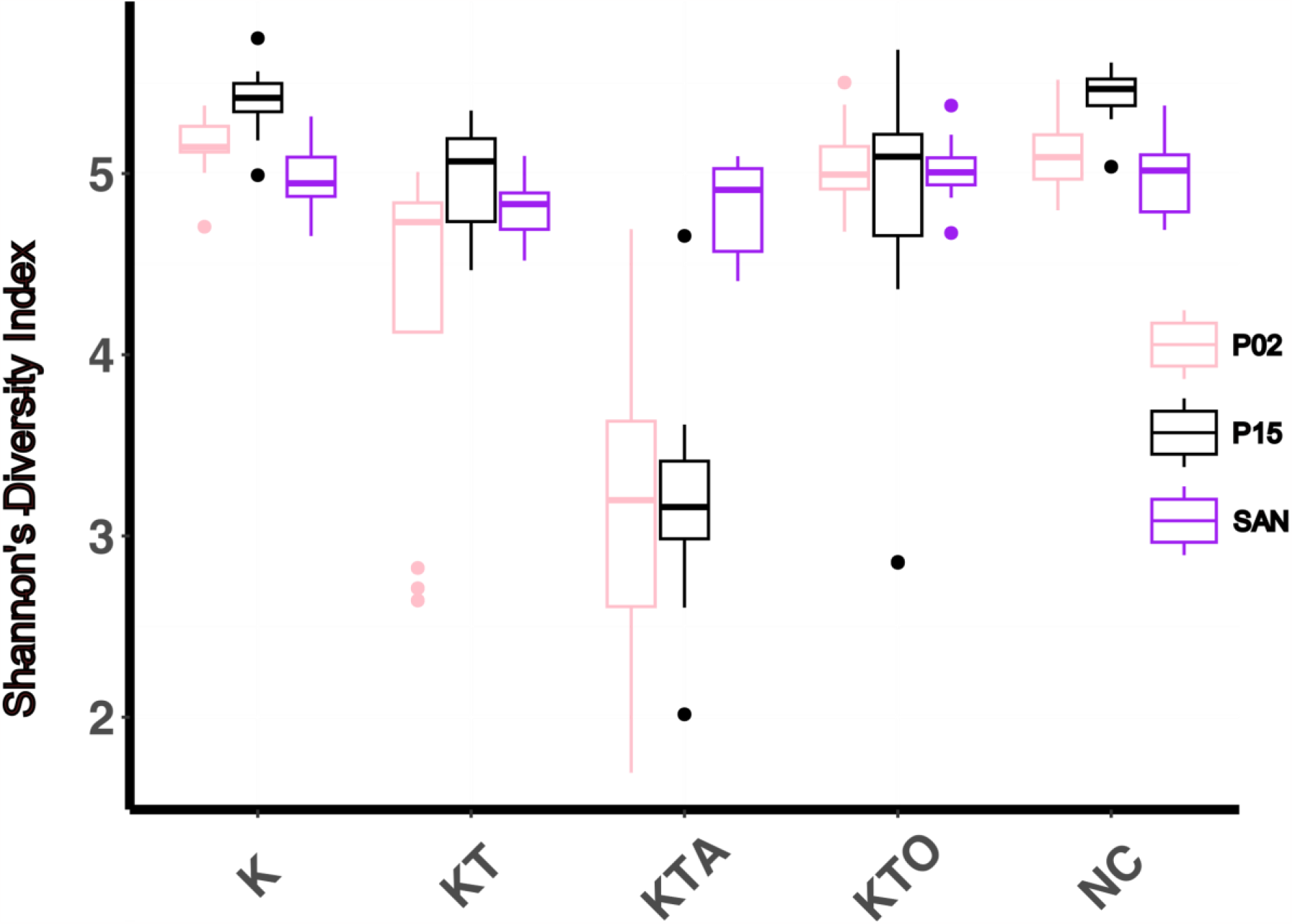
Shannon’s Diversity Index of samples from Experiment 1, summarized by chemical amendment on the X axis and grouped by soil type.

To probe whether the impact of chemical exposure could be detected at lower concentrations of chemical amendments, we varied the concentrations of each of the PUREX chemicals that the soils were exposed to and identified biomarker amplicon sequencing variants (ASVs) for chemical exposure, combining data over a series of separate experiments (Tables S1-S3). We specifically looked for ASVs with an increase in mean relative abundance from each experiment individually, and when data from all 3 experiments were pooled. Given these criteria, we identified 8 biomarker ASVs all of which were identified as biomarkers for the KT condition, whose abundance increased significantly in samples exposed to KT compared to other conditions. No biomarkers for K, KTO, or KTA were identified. However, the 8 ASVs from the KT condition typically also increased in abundance in the KTO condition, though the increase was not as great as in KT (Figure 3, Table 2). Three of these ASVs were in *Rhizobiaceae* family while the remaining 5 were in *Pseudomonas* genus. ASV7 and ASV15, both in *Rhizobiaceae*, were the highest relative abundance biomarkers (Table 2).

**Table 2.** Mean relative abundance for each of the 8 biomarker ASVs in each of the 5 conditions, and the fold change over negative control samples for KT and KTO conditions. The most specific assigned taxonomy for each ASV is shown.

| BiomarkerASV | NC<sup>1</sup> | K | KT | KTO | KTA | KT Fold Change | KTO Fold Change | Taxonomy |
| --- | --- | --- | --- | --- | --- | --- | --- | --- |
| ASV7 | 0.18 | 0.40 | 3.15 | 1.74 | 0.75 | 17.07 | 9.43 | Phyllobacterium<sup>2</sup> |
| ASV15 | 0.17 | 0.37 | 2.63 | 1.79 | 0.77 | 15.07 | 10.25 | Allorhizobium-Neorhizobium-Pararhizobium-Rhizobium |
| ASV19 | 0.18 | 0.46 | 2.39 | 2.05 | 0.53 | 13.36 | 11.50 | Pseudomonas |
| ASV27 | 0.18 | 0.41 | 2.14 | 1.79 | 0.38 | 12.11 | 10.09 | Pseudomonas |
| ASV35 | 0.09 | 0.24 | 1.58 | 1.17 | 0.60 | 17.89 | 13.22 | Pseudomonas |
| ASV54 | 0.20 | 0.19 | 0.81 | 0.83 | 0.16 | 4.01 | 4.12 | Pseudomonas |
| ASV62 | 0.03 | 0.12 | 0.92 | 0.44 | 0.23 | 31.37 | 15.14 | Rhizobiaceae |
| ASV94 | 0.02 | 0.07 | 0.76 | 0.64 | 0.18 | 41.52 | 34.93 | Pseudomonas |

<sup>1</sup>Negative control, no chemical added.
<sup>2</sup>Phyllobacterium is classified as a member of the *Rhizobiaceae* by SILVA (65) and GTDB (66), but as a member of the *Phyllobacteraceae* by NCBI taxonomy (67).

**Figure 3.**
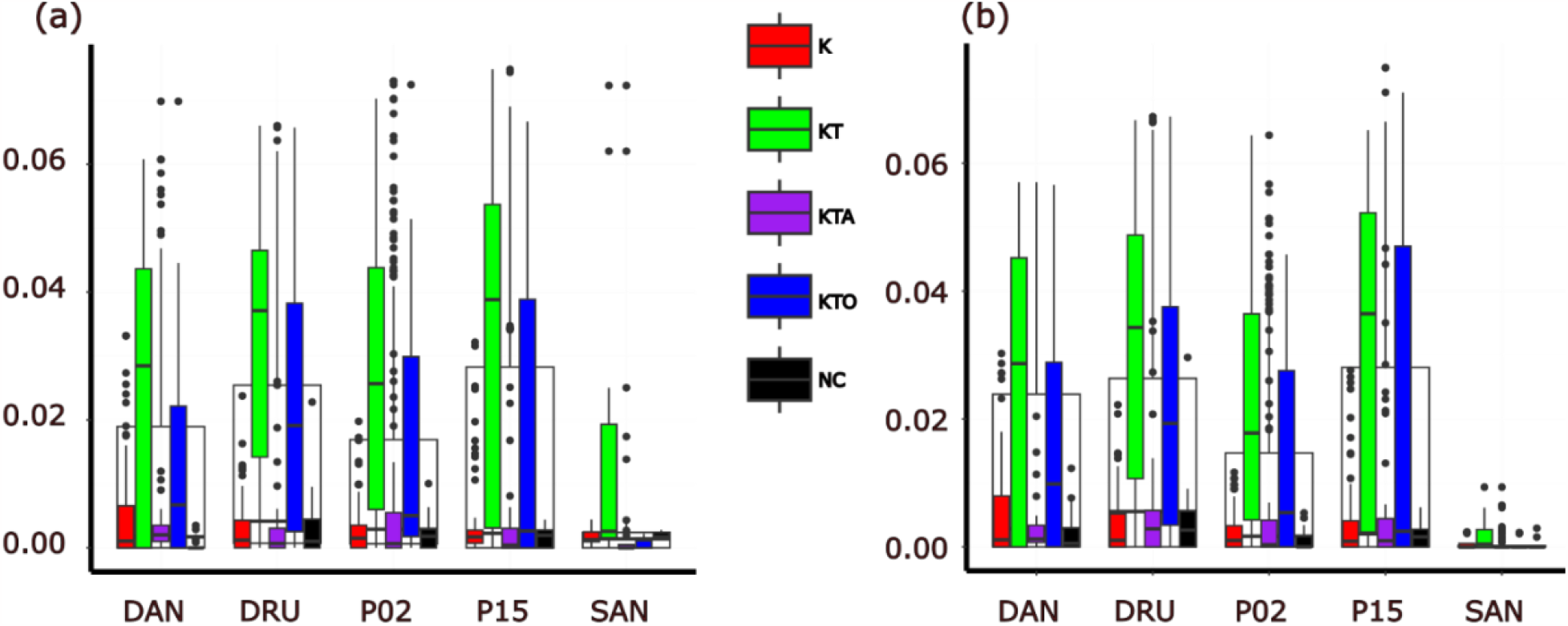
Relative abundance of ASV7 *Phyllobacterium* (a) and ASV15 (*Allorhizobium-Neorhizobium-Pararhizobium-Rhizobium*) (b), the two most consistent biomarker taxa. Samples from all 3 experiments shown summarized by soil type on X-axis and then clustered by chemical amendment. Black and white box and whisker plots behind colored box and whisker plots depict the overall sample values per soil type before further grouping into chemical conditions.

Using biomarker ASVs that were 20 fold more sensitive than whole community responses, we probed the lower limits of detection for chemical exposure and duration of response (Figure 4, Figure S1). The response of biomarker ASVs was weaker in sandy soils (SAN) (Figure 3). Biomarker ASVs also respond to KTO in some but not all soil types. For both KT and KTO conditions, biomarker response begins at a concentration of 5 μl/g at 7 days post exposure (Figure 4). For KT exposed samples responses persisted to 100 μl/g and out to 100 days post exposure, while responses to KTO were less durable. Most dramatically, KTO exposed samples had biomarker ASV abundances indistinguishable from negative controls at a concentration of 100 μl/g, while KT exposed samples had their strongest response at this concentration (Figure 4). This differing response is likely due to the drastically different pH between KT and KTO, which at higher concentrations may modify overall soil pH.

**Figure 4.**
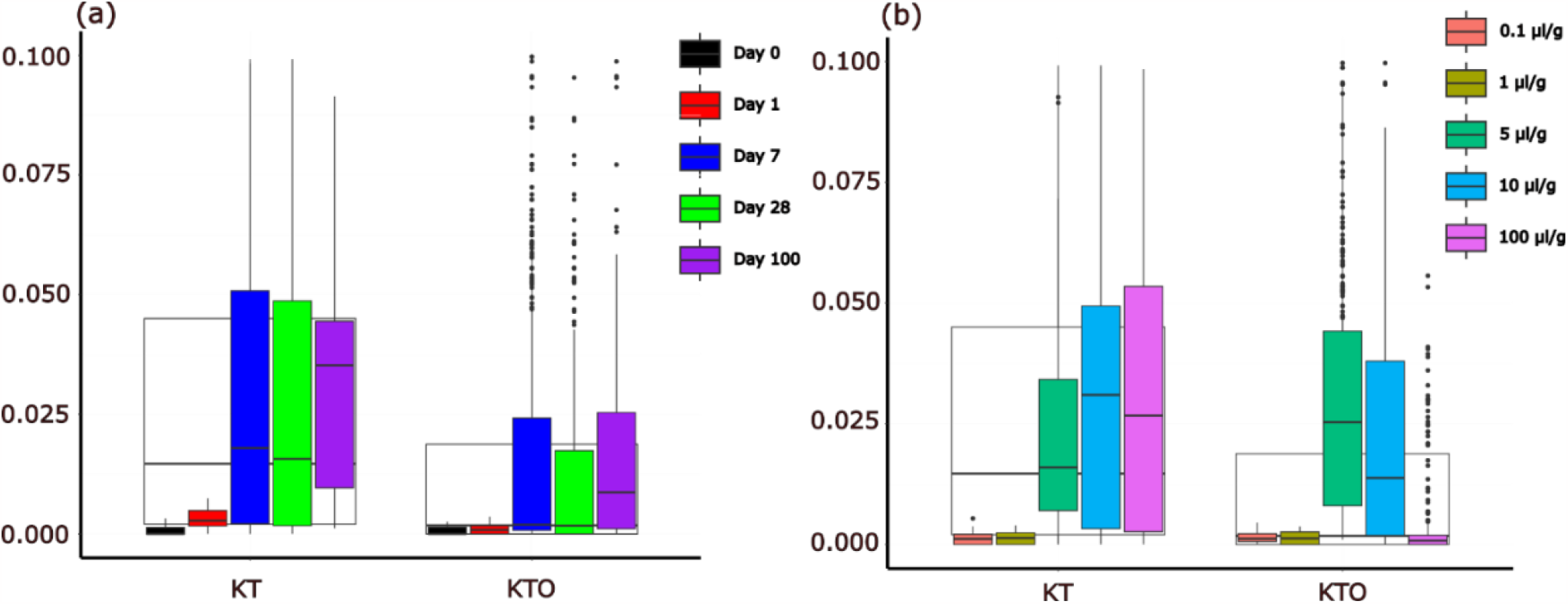
Relative abundance of ASV7 (*Phyllobacterium*) from all experiments, showing only samples exposed to KT or KTO conditions. (a) data are subset by the days post exposure of each sample per chemical amendment. (b) data are subset by the concentration of chemical amendment. Black and white box and whisker plots behind colored box and whisker plots depict overall sample values per chemical condition before further grouping into day (a) or concentration (b).

### Phosphotriesterase genes found in biomarker MAGs

We sought to reconstruct genomes that corresponded to the organisms represented by the 8 ASVs that increased in response to KT. We sequenced 96 shotgun metagenomic samples, generating 1,838 high quality metagenome assembled genomes (MAGs), 257 when dereplicated at a 95% ANI threshold. Using relative abundance from metagenomic read mapping, we identified 34 biomarker MAGs for KT exposed samples (Table S4). None of these 34 MAGs contained assembled 16S rRNA genes, so they could not be directly linked to the corresponding biomarker ASVs, although 15 of these MAGs fell into either family *Rhizobiaceae* or *Pseudomonadaceae*.

Since responses to KT were strongest and responses to K alone were not observed, we focused on tributyl phosphate as the causative agent for the growth of biomarker taxa. Using the 34 biomarker MAGs, we looked for genes that might be responsible for the degradation of tributyl phosphate. Microbially mediated degradation of tributyl is not well understood, but some known tributyl phosphate degraders have sequenced genomes. Using the genomes of *Sphingomonas* sp. RSMS (22) and *Rhodopseudomonas palustris* CGA009 (23), we performed a pan-genome analysis to determine if genes shared by biomarker MAGs and these two references genomes could be involved in the tributyl phosphate degradation pathway. We found no homology between any genes in our biomarker MAGs and CYP201A2 from *R. palustris* (24) or *aps* from *Sphingomonas* sp. RSMS (25). Therefore, we focused on phosphoesterase genes which may be involved in degrading tributyl phosphate to dibutyl phosphate (phosphotriesterases), dibutyl phosphate to monobutyl phosphate (phosphodiesterases) and monobutyl phosphate to butanol and inorganic phosphate (phosphomonoesterase or acid phosphatase) (25,26). Potential phosphtriesterases were identified, and their distribution across MAGs supports their role in promoting growth in response to tributyl phosphate. Searching for gene clusters in MAGs annotated as the above described phosphoesterases, we found more than 1,000 phosphodiestereases, at least one present in every biomarker MAG and also every one of the remaining 223 MAGs. We found no phosphomonesterases, though acid phosphatases are found in abundance, and the total number of acid phosphatase genes appears higher in many biomarker MAGs than on average amongst all MAGs (Table S4). We identified only 3 gene clusters annotated as phosphotriesterases-all which were found only in biomarker *Rhizobiaceae* MAGs.

Taking the sequences of these gene clusters annotated as phosphotriesterases, we quantified their abundance in 60 of the short read samples. We found that the relative copy number of phosphotriesterase genes in samples exposed to KT or KTO conditions were higher than in the control samples (Figure 5, Table 3). Phosphotriesterase copies per genome were on average 0.09 in negative control samples, 0.33 in KTO samples and 0.39 in KT samples (Pairwise Wilcoxon Rank Sum, KT vs. NC p= 4.8e-09, KTO vs. NC p= 1.3e-08). Samples exposed to 100 μl/g, 10 μl/g and 5 μl/g conditions were significantly higher than control samples for both KT and KTO (Table 3). Similar to biomarker ASVs and biomarker MAGs, phosphotriesterase copies per genome increased with increasing chemical concentration for KT exposed samples but not KTO exposed samples. For 5 μl/g,10 μl/g and 100 μl/g KT samples had 0.33, 0.42 and 0.43 phosphotriesterase copies per genome respectively. Meanwhile, for 5 μl/g,10 μl/g and 100 μl/g KTO samples had 0.34, 0.44 and 0.20 phosphotriesterase copies per genome respectively (Figure 5).

**Table 3:** Table of Wilcox rank test performed on values of phosphotriesterase genes per genome. * < 0.05, ** < 0.01, *** < 0.001, **** < 0.0001.

| Factor | Chi Squared | Degrees of Freedom | p value |
| --- | --- | --- | --- |
| **Chemical** | 29.352 | 2 | **** |
| KT vs NC |  |  | **** |
| KTO vs NC |  |  | **** |
| **Soil** | 7.562 | 1 | ** |
| **Concentration** | 32.516 | 3 | **** |
| 100 µl/g vs NC |  |  | **** |
| 10 µl/g vs NC |  |  | **** |
| 5 µl/g vs NC |  |  | **** |
| **Day** | 8.5857 | 2 | * |
| Day 28 vs Day 0 |  |  | * |
| Day 7 vs Day 0 |  |  | * |

**Figure 5:**
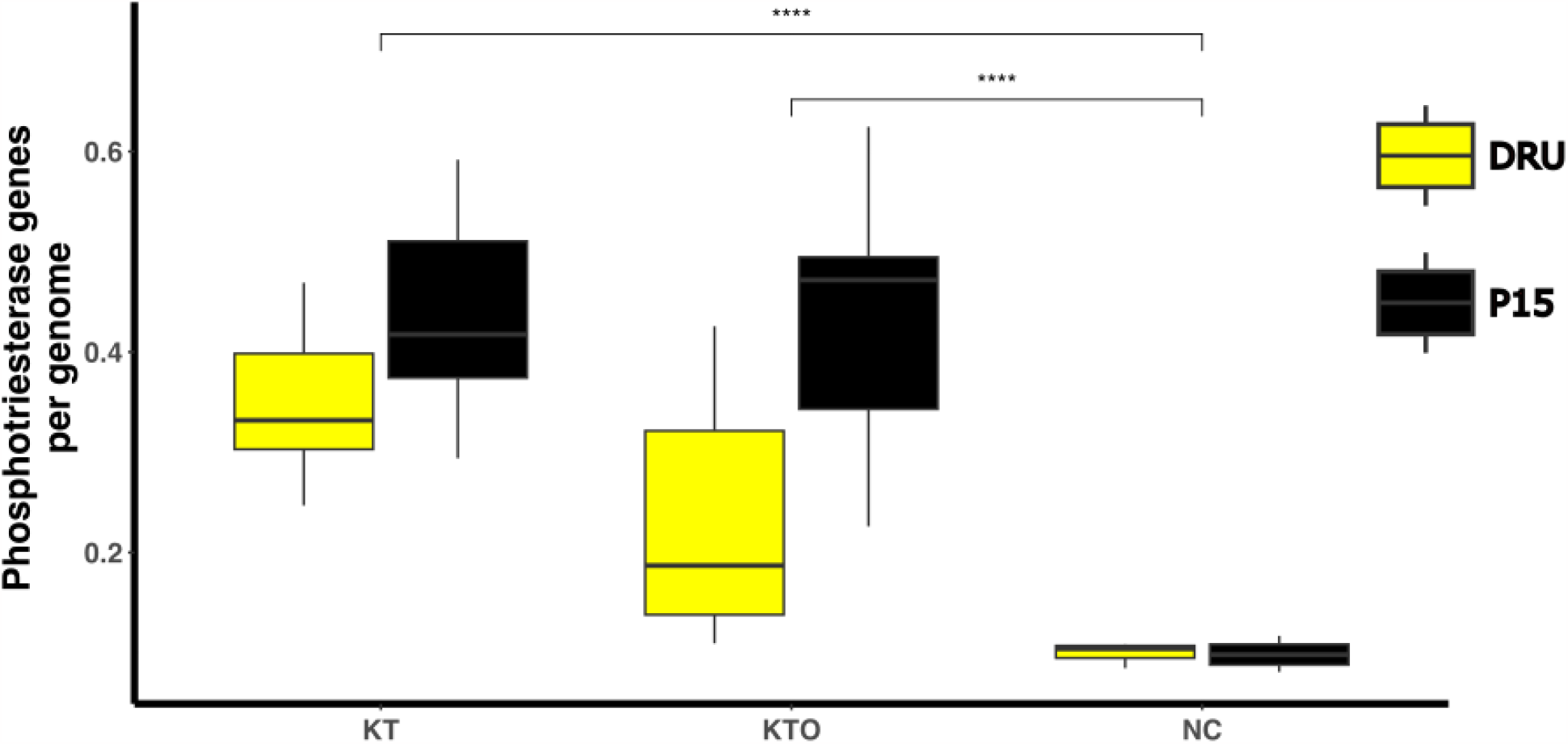
Phosphotriesterase genes per genome in metagenomic samples collected from Experiment 2. Summarized by condition on the X axis and then grouped by soil type. Statistical significance between groups (KT vs. NC, KTO vs. NC) indicated above boxes. * p < 0.05, ** p < 0.01, *** p < 0.001.

**Figure S1:**
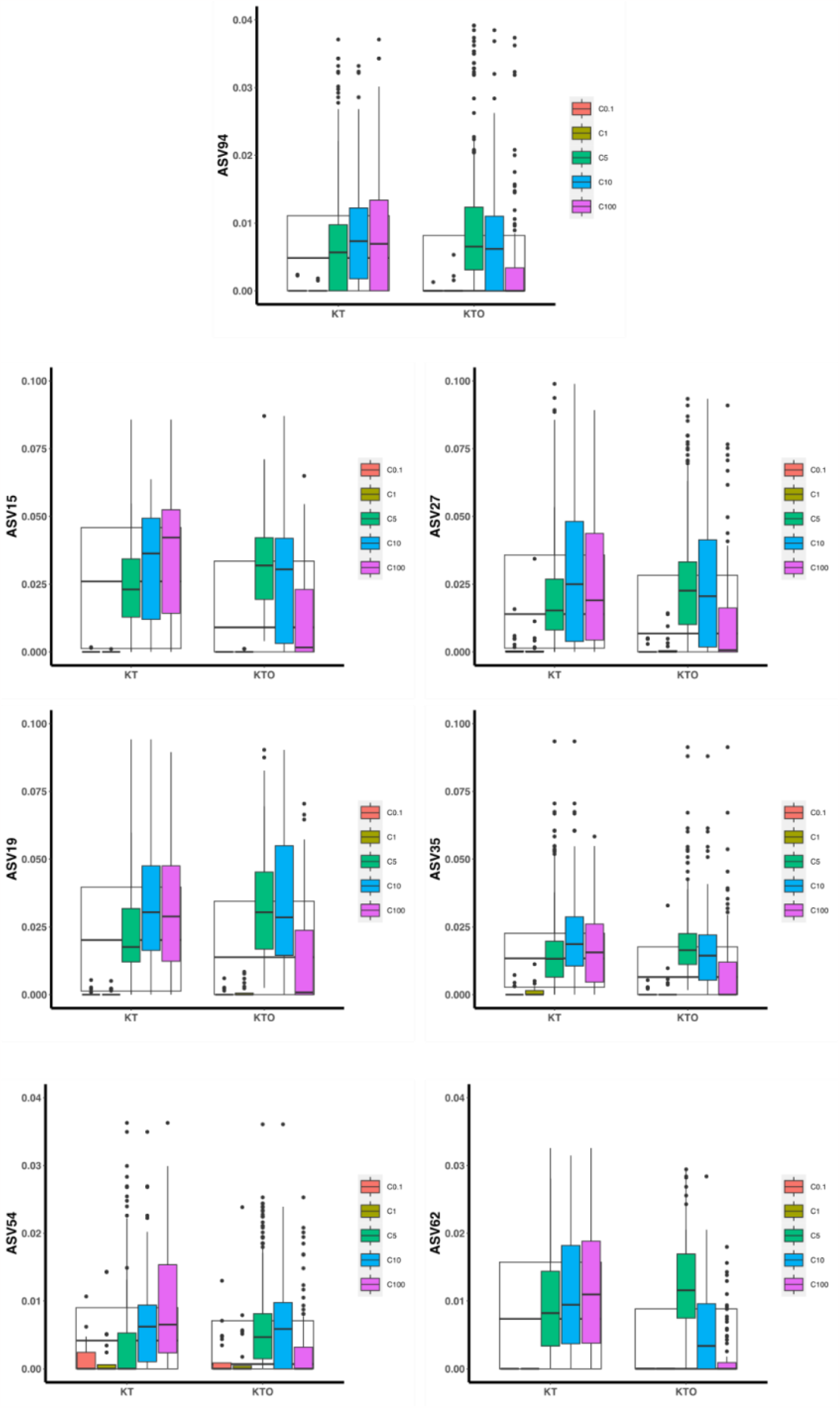
Relative abundance of biomarker ASVs, showing only samples exposed to KT or KTO conditions. Subset by concentration of chemical amendment within chemical amendment. ASV7 is shown in Figure 4.

**Table S1:** Experimental Conditions for Experiment 1. Soil Types defined in Table 1. Only a single concentration (100 μl/g) was used for this experiment. All conditions were carried out in triplicate.

| Soil Type | Chemical Amendment | Sample Times (Days) |
| --- | --- | --- |
| P02 | NC<br>K<br>KT<br>KTO<br>KTA | D0<br>D1<br>D7<br>D28 |
| P15 | NC<br>K<br>KT<br>KTO<br>KTA | D0<br>D1<br>D7<br>D28 |
| SAN | NC<br>K<br>KT<br>KTO<br>KTA | D0<br>D1<br>D7<br>D28 |

**Table S2:** Conditions for Experiment 2. Soil types are defined in Table 1. All conditions were carried out in triplicate except PC condition.

| Soil Type | Chemical Amendment | Chemical Concentration | Sample Times (Days) |
| --- | --- | --- | --- |
| P02 | NC<br>K<br>KT<br>KTO | C1 (0.1µl/g)<br>C2 (1 µl/g)<br>C3 (10 µl/g)<br>PC (100 µl/g) | D0<br>D7<br>D28 |
| P15 | NC<br>K<br>KT<br>KTO | C1 (0.1µl/g)<br>C2 (1 µl/g)<br>C3 (10 µl/g)<br>PC (100 µl/g) | D0<br>D7<br>D28 |
| DRU | NC<br>K<br>KT<br>KTO | C1 (0.1µl/g)<br>C2 (1 µl/g)<br>C3 (10 µl/g)<br>PC (100 µl/g) | D0<br>D7<br>D28 |
| DAN | NC<br>K<br>KT<br>KTO | C1 (0.1µl/g)<br>C2 (1 µl/g)<br>C3 (10 µl/g)<br>PC (100 µl/g) | D0<br>D7<br>D28 |

**Table S3:** Conditions for Experiment 3. Soil types are defined in Table 1. All conditions were carried out in triplicate.

| Soil Type | Chemical Amendment | Chemical Concentration | Sample Times (Days) |
| --- | --- | --- | --- |
| P02 | NC<br>K<br>KT<br>KTO<br>KTA | C5 (5 µl/g)<br>C10 (10 µl/g)<br>C100 (100 µl/g) | D0<br>D7<br>D28<br>D100 |
| P15 | NC<br>K<br>KT<br>KTO<br>KTA | C5 (5 µl/g)<br>C10 (10 µl/g)<br>C100 (100 µl/g) | D0<br>D7<br>D28<br>D100 |
| DRU | NC<br>K<br>KT<br>KTO<br>KTA | C5 (5 µl/g)<br>C10 (10 µl/g)<br>C100 (100 µl/g) | D0<br>D7<br>D28<br>D100 |
| DAN | NC<br>K<br>KT<br>KTO<br>KTA | C5 (5 µl/g)<br>C10 (10 µl/g)<br>C100 (100 µl/g) | D0<br>D7<br>D28<br>D100 |

**Table S4:** Selected MAGs identified as differentially abundant in KT condition samples. MAGs identified as being in Rhizobiaceae or Pseudomonas are shown. Number of genome clusters identified as phosphotriesterases or acid phosphatases in comparative genome analysis are shown for each MAG.

| MAG | GTDB_tk | Genome size (Mb) | % Completion | % Contamination | Phosphotriesterase | Acid Phosphatase |
| --- | --- | --- | --- | --- | --- | --- |
| P15_KTO_C10_R2_D28_E3_bin.19 | Rhizobium sp900470965 | 5.83 | 89.66 | 2.83 | 1 | 1 |
| P15_KT_PC_D28_E2_bin.10 | Rhizobium sp000799845 | 6.50 | 99.87 | 0.62 | 1 | 1 |
| P15_KTO_C10_R2_D7_E3_bin.15 | Rhizobium leguminosarum | 5.23 | 89.86 | 3.93 | 2 | 1 |
| Drum_KTO_C100_R1_D7_E3_bin.14 | Rhizobium altiplani | 5.63 | 97.49 | 0.67 | 1 | 2 |
| P15_KT_C100_R2_D28_E3_bin.19 | Pararhizobium sp000744565 | 5.61 | 83.61 | 3.63 | 0 | 0 |
| P15_KTO_C10_R2_D7_E3_bin.14 | Pararhizobium giardinii | 4.09 | 83.75 | 3.07 | 0 | 0 |
| P15_KT_C10_R2_D7_E3_bin.4 | Ensifer adhaerens A | 6.40 | 98.01 | 1.7 | 0 | 1 |
| P15_KT_C10_R1_D28_E3_bin.26 | Ensifer adhaerens | 4.76 | 77.17 | 3.61 | 1 | 2 |
| P2_KT_PC_D7_E2_bin.1 | Phyllobacterium | 5.81 | 99.91 | 0 | 1 | 2 |
| P15_KTO_C100_R2_D28_E3_bin.5 | Phyllobacterium | 5.95 | 98.62 | 0.87 | 1 | 3 |
| DRUM_KT_PC_D28_E2_bin.22 | Pseudomonas sp000282495 | 6.03 | 89.27 | 4.28 | 0 | 2 |
| P15_KT_PC_D28_E2_bin.14 | Pseudomonas E sp000282495 | 4.81 | 77.76 | 4.16 | 0 | 3 |
| P15_KTO_C10_R2_D7_E3_bin.12 | Pseudomonas E chlororaphis | 5.82 | 81.64 | 1.82 | 0 | 4 |
| P15_KT_C10_R2_D28_E3_bin.23 | Pseudomonas E | 4.23 | 90.08 | 3.76 | 0 | 1 |
| Drum_KTO_C5_R1_D28_E3_bin.6 | Pseudomonas E | 5.50 | 96.14 | 2 | 0 | 4 |

## Discussion

Our microcosm experiments demonstrated several key observations about the ways that the composition of soil microbial communities change in response to PUREX chemicals. Community-level changes occur when soils were amended with mixtures of kerosene/tributyl phosphate, kerosene/tributyl phosphate/nitric acid, but not kerosene alone. Responses of biomarker ASVs in *Rhizobiaceae* and *Pseudomonas* were more sensitive than whole community shifts and allow for recognition of exposure down to 5 μl/g. Genome reconstruction suggests an ability to degrade tributyl phosphate may drive the increased abundance of these biomarker ASVs. Quantification of phosphotriesterase genes suggest these genes may make a more generalizable biomarker than biomarker ASVs. Careful interpretation in light of our experimental design shows how our results may be applied to soil ecosystems and how they might respond to PUREX chemical exposure in a real-world contamination scenario.

One potential difficulty comparing our results to real world contamination events is that our microcosm experiments present a scenario in which PUREX chemicals are added to the soil but never removed, leading to a continual exposure over time. In real world exposures, rainwater would at some rate remove PUREX chemicals from soils (27). This would likely mean that the duration of the responses we observed in our experiments were greater than what would be present in a real-world contamination. Further, our method of exposure models a one-time release event, more consistent with catastrophic release rather than continuous exposure events (4). It is unclear if changes in community composition would be different in the same total concentration of PUREX chemicals were accumulated rather than were added in a single exposure event. Future mesocosm experiments may provide the ability to better bridge our results to real world scenarios.

Our results demonstrate that one soil type was less responsive to PUREX chemicals at both the community level and biomarker level. We found that microbiomes in sandy texture soils did not change in community composition in response to PUREX chemical amendments, and sandy soils produce muted biomarker ASV responses. The percentage of sand, which is associated with higher buffering capacity (28), was highest in the SAN soil. As we believe pH could drive changes at the community level especially to KTA condition, the greater buffering capacity may be why communities in sandy soil were not responsive.

Unlike KT, we observed decreasing biomarker ASVs in response to increasing concentration of KTO, which requires an understanding of the chemistry of kerosene, tributyl phosphate, and nitric acid mixtures. Dramatic responses of soil microbial communities to the aqueous phase of kerosene, tributyl phosphate and nitric acid (KTA) are likely due to the extremely low pH of this mixture. Given that the KTO chemical amendment was formed through phase separation from KTA, the pH and chemistry of KTO is clearly distinct from KT. Approximately 10% of the nitric acid from the initial aqueous phase is retained in the organic phase (29), and given our use of 4M nitric acid, retaining 0.4 M nitric acid has dramatic effects on the pH of KTO mixtures. A lower concentration of tributyl phosphate in KTO as compared to KT may also play a role, as hydrolysis of tributyl phosphate can occur in the KTO mixture (30). However, rates of acid hydrolysis of tributyl phosphate in the organic layer have been observed at approximately 1% of tributyl phosphate over 30 days even at much higher temperatures than used here (31). As a result, differing responses of soil communities to KT and KTO at the same concentration condition is likely a result of pH differences. And the loss of biomarker ASV response at 100 μl/g KTO is likely due to the higher total volume of mixture imparting a greater change on the total soil pH.

Our results suggest that the most pronounced and consistent biomarker responses occur in the KT condition. Given a lack of response to K alone, this suggests addition of tributyl phosphate to soils was the causative agent for growth of biomarker ASVs. Microbial degraders of tributyl phosphate are known, and some can degrade tributyl phosphate to butanol and phosphate, using tributyl phosphate as their sole source of carbon and phosphorus (32). Microbial degraders of tributyl phosphate are taxonomically diverse and found in *Rhodopseudomonas* (33), *Pseudomonas* (34), *Klebsiella* (35), and *Sphingomonas* (32). Members of *Rhizobiaceae* are not known to degrade tributyl phosphate, but have been shown to degrade similar organophosphates such as phorate and fenitrothion (19,36).

A genetic pathway for the degradation of tributyl phosphate is not complete for any tributyl phosphate degrading microbe. It is best characterized in *Sphingomonas* sp. RSMS (25,26), which is clearly distinct from the incomplete pathway characterization for *Rhodopseudomonas palustris* (24). As a result, the phosphostriesterase genes we observed as potentially involved with tributyl phosphate degradation may represent a novel pathway. We found that sequences with high amino acid identity to the phosphotriesterases discussed in this paper were present in numerous references such as *Rhizobium altiplani* BR 10423 (37) and *Neorhizobium petrolearium* DSM 26482 (38). As such, exploration of the tributyl phosphate degrading capabilities of these isolates could further confirm the role of this phosphotriesterase in tributyl phosphate degradation.

## Conclusions

Our work suggests that of PUREX chemicals, tributyl phosphate and nitric acid have the capacity to cause consistent changes in soil microbial community composition. Tributyl phosphate encourages the growth of *Rhizobiaceae* and *Pseudomonas*, while nitric acid broadly decreases diversity of communities, likely due to the toxic effects of extremely low pH. On the other hand, we did not observe detectable changes in microbial communities following kerosene exposure. Estimates of emissions from normal operations of a PUREX reprocessing tank are 0.0208 g/s of tributyl phosphate over a 5-day production cycle (4). Given that our 100 μl/g concentration condition contained approximately 0.029 g of tributyl phosphate per gram of soil, our experimental conditions should roughly model release scenarios during routine operations of nuclear materials reprocessing facilities. As a result, our identified biomarker taxa and genes may serve as tools to identify releases from PUREX reprocessing tanks.

## Methods

### Chemical Selection

Chemical conditions were selected based on their use in PUREX, their identity as semi volatile, and their potential for environmental impact (4). Ratios and concentrations of chemicals were based on those used in traditional PUREX chemistry flow sheets (39,40). Kerosene (Sigma Aldrich 60710), tributyl phosphate (Sigma Aldrich 00675) and nitric acid (Fisher Scientific A467-250) were used. Mixtures for chemical conditions were made as follows. K condition was 300 ml kerosene. KT condition was 210ml Kerosene and 90ml tributyl phosphate (TBP). For the KTA and KTO conditions, nitric acid was diluted to 4M. Then, 350ml of 4M nitric acid was added to 350ml of 30% tributyl phosphate 70% kerosene mixture. This was vortexed for 30 seconds and then incubated until phase separation occurred. Aqueous phase (KTA) and organic phase (KTO) were then aspirated into separate containers.

### Soil Collection

Soil was collected from sites described in Table 1. Clay and soil organic matter are considered the most chemically reactive components of soil. The cation exchange capacity (CEC) of soil has a large influence on how soils interact with nutrients and chemicals. CEC is influenced by the % clay and the organic matter content, both of which impart a negative charge to soils. Soils representing several soil types were collected. By using soils with similar chemical and physical characteristics but different microbial communities, we will be able to determine if the chemical treatments impose characteristic and predictable effects on the microbial communities by looking for microbial community changes that are similar across the different communities. Selection of a variety of contrasting soil types/textures allows us to determine if there are universal effects of the PUREX contaminants on the microbial community. Soil was collected from the top 10 cm of topsoil using a shovel, avoiding plant crowns. After collection, soil was stored at 4 ° C for 1 week week before use. Insects, roots, sticks and other large materials were removed from soil before use. For repeated soil types, new soil was collected before each experiment. Soils texture and chemical analysis was performed by Waypoint Analytical (Champaign, Illinois).

### Experimental Setup

For Experiment 1, soil types P02, P15 and SAN were used. 100 μl/g of soil of each solution was added to soil using a pipette: Kerosene, 70% Kerosene / 30% TBP, 70% Kerosene / 30% TBP with 4M nitric acid (organic layer), 70% Kerosene / 30% TBP with 4M nitric acid (aqueous layer), DI water (negative control, NC). Each condition was carried out in triplicate. Post addition of chemical, microcosms were incubated at 20° C with 50% humidity and destructively sampled at 0-, 1-, 7-, and 28-days post exposure. DNA was extracted from 250 mg of soil.

For Experiment 2, soils P02, P15, DAN and DRU were used. 0.1 μl/g, 1.0 μl/g, 10 μl/g or 100 μl/g of each solution was added to soil: Kerosene, 70% Kerosene / 30% TBP, 70% Kerosene / 30% TBP with 4M nitric acid (organic layer), No addition (NC). Each condition was carried out in triplicate, except positive control (PC) 100 μl/g. 100 μl/g was added via pipette while other concentrations were dispersed via a glass nebulizer. 275 μl was placed in the nebulizer to achieve 0.1 μl/g, 812 μl was placed to achieve 1.0 μl/g, and 6182 μl was placed to achieve 10 μl/g. Post addition of chemical, microcosms were incubated at 20° C with 50% humidity and destructively sampled at 1-, 7-, and 28-days post exposure. DNA was extracted from 250 mg of soil.

For Experiment 3, soils P02, P15, DAN and DRU were used. 5 μl/g, 10 μl/g or 100 μl/g of each solution was added to soil: Kerosene, 70% Kerosene / 30% TBP, 70% Kerosene / 30% TBP with 4M nitric acid (organic layer), 70% Kerosene / 30% TBP with 4M nitric acid (aqueous layer), No addition (NC). Each condition was carried out in triplicate.100 μl/g was added via pipette while other conditions were dispersed via a glass nebulizer. 3,198 μl was placed in the nebulizer to achieve 5 μl/g and 6182 μl was placed to achieve 10 μl/g. Post addition of chemical, microcosms were incubated at 20° C with 50% humidity and destructively sampled at 7-, 28- and 100-days post exposure. DNA was extracted from 250 mg of soil.

### DNA Sequencing

DNA was extracted from soil using DNeasy PowerSoil Kit (Qiagen 47014). For 16S rRNA sequencing, amplification, library preparation and sequencing were carried out at the Argonne National Laboratory Environmental Sample Preparation and Sequencing Facility. V4 primers 515F (41) – 806R (42) were used for amplification. DNA sequencing was carried out on an Illumina Miseq. For shotgun metagenomic sequencing, library preparation and DNA sequencing was carried out at the Roy J Carver Biotechnology Center at the University of Illinois at Urbana-Champaign. Library preparation was carried out using Hyper Library Construction kit from Kapa Biosystems, and DNA sequencing was performed on an Illumina NovaSeq 6000 S4 Flow Cell, 2x 150 bp reads. Shotgun metagenomic sequencing was carried out for 96 samples: 36 samples from Experiment 2 and 60 samples from Experiment 3.

### Computational Analysis

16S rRNA samples were analyzed in DADA2 (43) following the standard tutorial workflow in R (44). Exact commands used are described in code repository (https://github.com/jpod1010/microbiomestoassessproductionfacilities). Ordination and alpha diversity calculation were carried out in phyloseq (45). LEfSE biomarker discovery (46) and DEseq2 biomarker discovery (47) were carried out in microbiomeMarker (48). Plots were generated using ggplot2 (49), phyloseq or R package microbiome. Biomarker designation was assigned based to ASVs that were identified by DEseq2 and LeFSe in all experiments independently and when all experimental data was combined.

Shotgun metagenomic samples were quality controlled, processed and analyzed using custom scripts. Each shotgun metagenomic sample was qtrimmed at trimq=30 qtrim=r with bbduk (50). Digital normalization was carried out using bbnorm with target=40 mindepth=2. Assembly with megahit was performed on quality trimmed and digitally normalized sequences using –presets meta-large. For the 36 samples in Experiment 1, bwa (51) was used through metaWRAP (52) was used to map short reads from all of the 36 quality-controlled short read samples onto each of the 36 assemblies, and metabat2 (53), maxbin2 (54) and CONCOCT (55) were used to bin based on those coverages. MetaWRAP bin_refinement was used to consolidate these three bin sets based on CheckM 1.0.12 (56) and in all this produced 36 sets of bins. This same process was carried out for the 60 samples taken from Experiment 2, mapping quality controlled short reads from Experiment 2 to assemblies from Experiment 2. Together this generated 96 sets of bins from experiments 1 and 2. We then used dRep (57) to dereplicate MAGs at 95% identity using - sa 0.95. We assigned taxonomy to these MAGs using GTDB-tk (58). We then used metaWRAP quant_bins to quantify the abundances of these MAGs in short reads from experiment 1 and experiment 2 separately. For Experiment 2, these MAG abundance values were imported into phyloseq and microbiomeMarker was used to run LeFSe and Deseq2 on these abundances to identify biomarker MAGs. These biomarker MAGs, along with Sphingomonas sp. RSMS (GCF_005406185.3) and Rhodopseudomonas palustris CGA009 (GCF_026625025.1), were imported into Anvi’o (59), annotated using KEGG (60), Pfam (61), and COG (62), and a pan-genome analysis was carried out. Genes of interest annotated as phosphotriesterases were exported and those sequences were searched against quality controlled short reads from all 96 samples using DIAMOND BLASTx (63). MicrobeCensus (64) was run on all 96 quality controlled short reads to identify the number of genome equivalents present in each sample, and this was used to normalize the number of reads identified as phosphotriesterases in each sample. These normalized values were imported into R and ggplot2 was used to generate figures and Wilcox Signed Rank tests were carried out using stats package.

## Data Availability

Raw 16S rRNA sequence files, raw shotgun metagenome sequence files, and genome assemblies are available under BioProject PRJNA1000596. Code used to analyze 16S data, shotgun metagenome data, and generate figures is available on GitHub (https://github.com/jpod1010/microbiomestoassessproductionfacilities).

## Acknowledgements

Funding was provided by the National Nuclear Security Agency, Office of Nonproliferation Research and Development (NA-22), through grant AN-19-Microbiome-PD3RA. The authors would like to thank Ellie Hailmann, Rachel Waltermire, Ingrid Holstrom, and Kristin Ellis for their assistance in carrying out laboratory work.

## Author contributions

All authors contributed to experimental design. S.F carried out experiments. J.C.P carried out bioinformatic analysis. J.C.P wrote manuscript. All authors contributed to revision and review of manuscript.

